# The sequence of a male-specific genome region containing the sex determination switch in *Aedes aegypti*

**DOI:** 10.1101/122804

**Authors:** Joe Turner, Ritesh Krishna, Arjen E. Van’t Hof, Elizabeth R. Sutton, Kelly Matzen, Alistair C. Darby

**Affiliations:** Centre for Genomic Research, Institute of Integrative Biology, University of Liverpool, 9 Crown Street, Liverpool, L69 7ZB, UK.; Oxitec Ltd., 71 Innovation Drive, Milton Park, Abingdon, OX14 4RQ, UK.; Department of Zoology, University of Oxford, South Parks Road, Oxford, OX1 3PS, UK.

## Abstract

*Aedes aegypti* is the principal vector of several important arboviruses. Among the methods of vector control to limit transmission of disease are genetic strategies that involve the release of sterile or genetically modified non-biting males (Alphey 2014), which has generated interest in manipulating mosquito sex ratios (Gilles *et al.* 2014; Adelman and Tu 2016). Sex determination in *Ae. aegypti* is controlled by a non-recombining Y chromosome-like region called the M locus (Craig *et al*. 1960), yet characterisation of this locus has been thwarted by the repetitive nature of the genome (Hall *et al.* 2015). In 2015, an M locus gene named Nix was identified that displays the qualities of a sex determination switch (Hall *et al.* 2015). With the use of a whole-genome BAC library, we amplified and sequenced a ~200kb region containing this male-determining gene. In this study, we show that Nix is comprised of two exons separated by a 99kb intron, making it an unusually large gene. The intron sequence is highly repetitive and exhibits features in common with old Y chromosomes, and we speculate that the lack of recombination at the M locus has allowed the expansion of repeats in a manner characteristic of a sex-limited chromosome, in accordance with proposed models of sex chromosome evolution in insects.

---

At least 2.5 billion people live in areas where they are at risk of dengue transmission from mosquitoes, principally *Ae. aegypti,* with an estimated 390 million infections per year (Laughlin *et al.* 2012; Bhatt *et al.* 2013). Recently, the emergence of chikungunya and Zika viruses further highlights the public health importance of *Ae. aegypti* (Musso *et al.* 2015; Fauci and Morens 2016). Future mosquito control strategies may incorporate genetic techniques such as the sustained release of sterile or transgenic “self-limiting” mosquitoes (Alphey et al., 2013; WHO: https://goo.gl/FRqJ0d). Given that only female mosquitoes bite and spread disease, there has been substantial interest in manipulating mosquito sex determination using these genetic techniques and others, including gene drive (Adelman and Tu 2016; Hoang *et al.* 2016). Therefore, elucidating the genetic basis for sex determination could, for instance, facilitate production of male-only cohorts for release, or allow transformation of mosquitoes with sex-specific “self-limiting” gene cassettes.

Sex determination in insects is variable, and generally not well understood outside of model species (Charlesworth and Mank 2010). Unlike the malaria mosquito *Anopheles gambiae* and *Drosophila* species, *Ae. aegypti* does not have heteromorphic (XY) sex chromosomes (Craig *et al.* 1960). Instead, the male phenotype is determined by a non-recombining M locus on one copy of autosome 1 (Newton *et al.* 1978; Clements 1992; Toups and Hahn 2010). This locus is poorly characterised because its highly repetitive nature has confounded attempts to study it based on the existing genome assembly (Hall *et al.* 2015). The 1,376Mb *Ae. aegypti* genome was assembled from Sanger sequencing reads in 2007 (Nene *et al.* 2007), which are commonly not long enough to span the repetitive transposable elements that comprise a large proportion of the genome (Koren and Phillippy 2015). Consequently, the current assembly is still relatively low quality (Severson and Behura 2012). Furthermore, the fact that both male and female genomic DNA was used for genome sequencing reduces the expected coverage of the M locus to one quarter of the autosome 1 sequences, further obscuring candidate M locus sequences (Hall *et al.* 2014).

Recently, a team of researchers was nevertheless able to identify *Nix,* a gene with male-specific, early embryonic expression. Knockout of *Nix* using CRISPR/Cas9 results in morphological feminisation of male mosquitoes along with feminisation of gene expression and female splice forms of the conserved sex-regulating genes *doublesex (dsx)* and *fruitless (fru),* strongly indicating that *Nix* is the upstream regulator of sexual differentiation (Hall *et al.* 2015). The translated *Nix* protein contains two RNA recognition motifs and is hypothesised to be a splicing factor, acting either directly on *dsx* and *fru* or on currently unknown intermediates (Adelman and Tu 2016). A comparison of sexually dimorphic gene expression in different mosquito tissue types also detected male-specific transcripts of *Nix* (Matthews *et al.* 2016). An ortholog of *Nix* is present in *Ae. albopictus,* but it is not known if the two are functionally homologous (Chen *et al.* 2015).

To date, *Nix* has only been characterised as an mRNA transcript. To fully understand this gene’s role in sex determination and to utilise this knowledge for vector control, it is essential to decipher its genomic context. For this purpose, this study identifies and describes the region of the M-locus in which *Nix* is located.

Four BAC clones positive for *Nix* assembled into a single region of 207 kb with no gaps and a GC content of 40.2% (submitted to the NCBI as accession KY849907). The presence of the *Nix* gene in the assembled BACS was confirmed by BLASTN. The whole gene was present in tiled BACs, though not completely within individual BAC clones. Neither *Nix* nor the complete region could be found in the AaegL3 or Aag2 reference genome assemblies. While *Nix* was originally identified in the genome-sequenced Liverpool strain (Hall *et al.* 2015), PCR revealed that it is exclusively present in male genomic DNA from other geographically varied *Ae. aegypti* populations (Figure S1), further strengthening the evidence that it is wholly present in the M locus.

The *Nix* gene was found to be made up of two exons with a single intron of 99 kb (Figure 1). Although large introns are not uncommon in *Ae. aegypti* (average intron length ~5000 bp)(Nene *et al.* 2007), this intron is at the extreme end of intron sizes observed (Figure S2), especially considering the small size of its protein coding regions (<1000 bp). The gene structure is confirmed by Illumina RNA-Seq data clearly showing reads spanning the intron between the two exons (Figure 1). RepeatMasker identified approximately 55% of the sequenced region as repetitive, and the intron region of *Nix* as 72% repetitive (Table S1).

**Figure 1:**
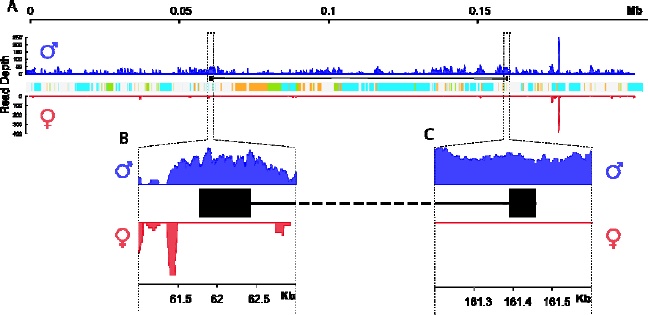
Structure and gene expression of the ~207 kb genomic region containing the *Nix* gene. *Nix* is shown as two black boxes representing the exons, joined by a black line representing the intron. Colours on the central track of **A** represent the classes of repetitive elements (orange: DNA transposons; cyan: Gypsy LTRs; green: Ty1/Copia LTRs). Blue histograms represent the coverage of RNA-Seq reads from male samples on the *y* axis; red histograms represent the coverage from female samples. **B** and **C** show enlargements of the first and second exons of *Nix* in the dotted regions in **A**, respectively.

The genomic data from our assembled M locus region show that *Nix* is approximately 100 kb in length – exceptionally long even for an insect, and one of the longest in the mosquito genome. This is particularly unusual because *Nix* is expressed in early embryonic development, before the onset of the syncytial blastoderm stage 3-4 hours after oviposition (Hall *et al.* 2015), during which time most active genes have very short introns, or lack them entirely. There is evidence of selection against intron presence in genes expressed in the early *Ae. aegypti* zygote (Biedler *et al.* 2012). In *Drosophila,* the majority of early-expressed genes have small introns and encode small proteins, suggesting that selection has favoured high transcript turnover during early embryonic development due to the requirement for short cell cycles and rapid division (Artieri and Fraser 2014). It might therefore be expected that selection would limit the *Nix* intron’s expansion to preserve efficient transcription in the zygote.

One possible explanation is the expansion of repetitive DNA. The RepeatMasker results reveal that the *Nix* region contains a high number of repetitive sequences, especially retrotransposons (Figure 1; Table S1). The M locus has accumulated repeats in between protein-coding DNA in a manner characteristic of a sex chromosome, which are prone to degeneration by Muller’s ratchet due to the lack of recombination (Muller 1964; Charlesworth 1991; Kaiser and Bachtrog 2010). For instance, repetitive sequences comprise almost the entire *Anopheles gambiae* Y chromosome, and these repetitive sequences show rapid evolutionary divergence (Hall *et al.* 2016). Similarly, genes on the *Drosophila* Y chromosome, such as those involved in spermatogenesis, have gigantic repetitive introns, sometimes in the megabase range, that consequently make them many times larger than typical autosomal genes (Carvalho *et al.* 2001; Bachtrog 2013).

It is therefore possible that the lack of recombination may pose constraints on the structure of the M locus, and in the absence of strong selection the *Nix* gene has degenerated outside the coding regions. Non-recombining sex loci such as the *Ae. aegypti* M locus may represent an evolutionary precursor to differentiated sex chromosomes, which are thought to emerge when sexually antagonistic alleles accumulate on either chromosome and favour reduced recombination between the two homologs, eventually leading to degeneration and loss of genes on the proto-Y (Charlesworth *et al.* 2005). Recent data appears to show that recombination is reduced along autosome 1 even outside of the M locus(Fontaine *et al.* 2016), while the fully differentiated *Anopheles* X and Y chromosomes still display some degree of recombination with each other (Hall *et al.* 2016). Thus, *Ae. aegypti* may be “further along” this evolutionary trajectory than previously assumed.

The *Ae. aegypti* M locus provides an intriguing example of the complexity of evolutionary forces acting on sex chromosomes, and further study of the locus will contribute to understanding the evolution of sex determination in insects and address general questions about the factors impacting gene and genome length. Importantly, these may also yield insights that can be applied to increase the efficiency of genetic strategies for vector control.

## Methods

### BAC library construction

A BAC library of insert size 130 kb was constructed (Amplicon Express, USA) for an estimated coverage of ~5x for autosomal regions (~2.5x for sex specific regions) from a DNA pool of approximately 50 sibling males. The male siblings were from one family from an Asian wild type laboratory strain after five generations of full-sib mating. Superpools and matrixpools were supplied to allow PCR based screening of the BAC library.

### BAC library screening, isolation and sequencing

The BAC library was PCR screened using primers (Nix1F 3’-TTGAGTCTGAAAAGTCTATGCAA-5’, Nix1R 3’-TCGCTCTTCCGTGGCATTTGA-5’, Nix2F 3’- ACGTAGTCGGCAACTCGAAG-5’, Nix2R 3’-CTGGGACAAATCGAACGGAA-5’) based on the complete coding sequence of *Nix* (GenBank accession number KF732822). The first primer set was also used to screen for *Nix* in the genomic DNA of six male and six female individuals each from two wildtype *Ae. aegypti* strains.

Screening of the library resulted in four positive clones - two for each primer pair. These BAC clones were propagated, extracted using a Maxiprep kit (Qiagen, UK), pooled before SMRTbell library preparation (PacBio, USA), and sequenced on a single SMRTcell using P6-C3 chemistry on the PacBio RS II platform (PacBio, USA).

### Data analysis

The sequence data was trimmed to remove vector sequences and adaptors prior to assembly with the CANU v1 assembler (Berlin *et al.* 2015), followed by sequence polishing with QUIVER.

BLASTN was used to assess the uniqueness of the assembled *Nix* region compared to the *Aedes aegypti* Liverpool reference genome AaegL3 and the newer Aag2 cell line assembly. Illumina data generated from male and female genomic DNA (accession numbers SRX290472 and SRX290470) and RNA (accession numbers SRX709698-SRX709703) were mapped to a combined reference containing the assembled *Nix* region added to the AaegL3 genome. DNA samples were mapped with BOWTIE 2.2.1 (using default parameters with–I 200 and-X 500) and RNA-Seq data with TOPHAT 2.1.1 version (using default parameters). RNA-Seq data was processed using the CUFFLINKS 2.2.1 pipeline to look for potential genes and male/female specific expression from the region.

Genes were predicted using AUGUSTUS and the *Aedes aegypti* model (Nene *et al.* 2007), repetitive regions described using REPEATMASKER 4.0.6 and the *Ae. aegypti* repeat database.

**Supplementary Information** is available in the online version of the paper.

## Author contributions

J.T., R.K. and A.E.v.H. contributed equally to this work. K.M. and A.C.D. designed the study and obtained funding, with contribution from J.T.; K.M. provided mosquito samples; E.R.S. and A.C.D. commissioned the BAC library construction; A. E. v. H. and J. T. screened the BAC library and extracted DNA; A. E. v. H. performed BAC scaffolding; A.C.D. oversaw sequencing and assembled the DNA sequence; R.K. performed the mapping and developed computational strategies for data analysis; J.T. performed the repeat masking; J.T. and A.C.D. wrote the paper, with contribution from A. E. v. H.; R.K. and A.C.D. produced the figures.

## Acknowledgments

This work was funded by BBSRC PhD training grant BB/M503460/1 (J.T. & A.C.D.) and a BBSRC grant BB/M001512/1 (K.M. & A.C.D.).

The PacBio sequencing was conducted at the Centre for Genomics Research, University of Liverpool with the assistance of Dr Margaret Hughes and Dr John Kenny.

We thank Dr Andrea Betancourt and Dr Ilik Saccheri for comments on the manuscript.

## Supplementary Information

**Figure S1:**
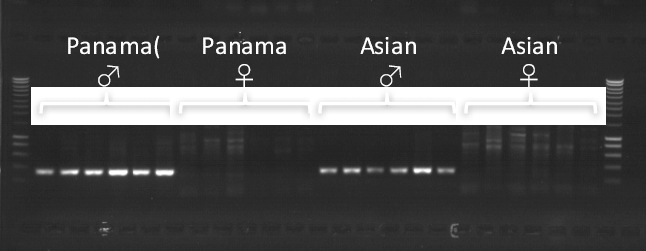
PCR screening of the M locus gene *Nix* in male and female DNA of wild type *Aedes aegypti* strains. Primers used were Nix1F (3’-TTGAGTCTGAAAAGTCTATGCAA-5’) and Nix1R (3’-TCGCTCTTCCGTGGCATTTGA-5’), targeting Nix exon 1.

**Figure S2:**
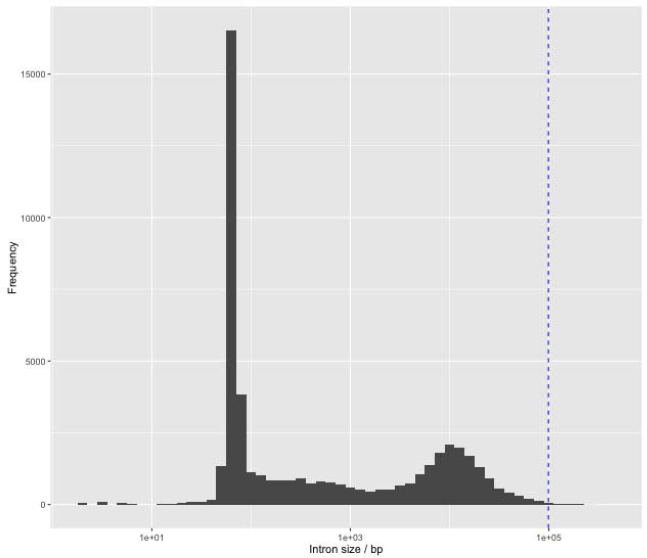
Figure S2: Intron size distribution in *Aedes aegypti* Liverpool reference genome AaegL3. Blue dashed line indicates the size of the *Nix* intron relative other introns. X axis is transformed by log_10._

**Table S1:** Types and abundance of repeats in the 207kb assembled M locus region and 99kb *Nix* intron, identified by RepeatMasker using the *Aedes aegypti* repeat library.

| Repeat Type | Entire region |  | *Nix* intron region |  |
| --- | --- | --- | --- | --- |
|  | Number of elements | Percentage of sequence | Number of elements | Percentage of sequence |
| **Retroelements** | 105 | 42.1% | 49 | 51.0% |
| **SINEs** | 8 | 0.81% | 5 | 1.11% |
| Penelope | 3 | 0.08% | 2 | 0.20% |
| **LINEs** | 24 | 5.43% | 6 | 6.85% |
| L2/CR1/Rex | 4 | 0.13% | 0 | 0% |
| R1/L0A/Jockey | 13 | 3.87% | 3 | 6.60% |
| RTE/Bov-B | 3 | 1.33% | 0 | 0% |
| L1/CIN4 | 1 | 0.02% | 1 | 0.05% |
| **LTR Elements** | 73 | 35.8% | 38 | 43.0% |
| BEL/Pao | 9 | 0.71% | 3 | 0.87% |
| Ty1/Copia | 16 | 11.3% | 14 | 19.2% |
| Gypsy/DIRS1 | 48 | 23.8% | 21 | 23.0% |
| **DNA transposons** | 97 | 11.7% | 69 | 20.1% |
| Tc1-IS630-Pogo | 11 | 3.87% | 11 | 9.04% |
| Other (Mirage, P-element, Transib) | 1 | 0.06% | 0 | 0% |
| **Unclassified** | 6 | 0.48% | 3 | 0.22% |
| **Small RNA** | 8 | 0.81% | 5 | 1.11% |
| **Satellites** | 1 | 0.75% | 0 | 0% |
| **Simple repeats** | 19 | 0.34% | 7 | 0.24% |
| **Low complexity** | 3 | 0.07% | 1 | 0.04% |
| **Total repeats** |  | 55.4% |  | 71.6% |

## References

Adelman Z. N., Tu Z., 2016 Control of mosquito-borne infectious diseases: sex and gene drive. Trends Parasitol. 32: 219–229.

Alphey L., McKemey A., Nimmo D., Neira Oviedo M., Lacroix R., et al., 2013 Genetic control of *Aedes* mosquitoes. Pathog. Glob. Health 107: 170–179.

Alphey L., 2014 Genetic control of mosquitoes. Annu. Rev. Entomol. 59: 205–224.

Artieri C. G., Fraser H. B., 2014 Transcript length mediates developmental timing of gene expression across *Drosophila*. Mol. Biol. Evol. 31: 2879–2889.

Bachtrog D., 2013 Y-chromosome evolution: emerging insights into processes of Y-chromosome degeneration. Nat. Rev. Genet. 14: 113–124.

Berlin K., Koren S., Chin C.-S., Drake J. P., Landolin J. M., et al., 2015 Assembling large genomes with single-molecule sequencing and locality-sensitive hashing. Nat. Biotechnol. 33: 623–630.

Bhatt S., Gething P. W., Brady O. J., Messina J. P., Farlow A. W., et al., 2013 The global distribution and burden of dengue. Nature 496: 504–507.

Biedler J. K., Hu W., Tae H., Tu Z., 2012 Identification of early zygotic genes in the yellow fever mosquito *Aedes aegypti* and discovery of a motif involved in early zygotic genome activation. PLoS One 7: e33933.

Carvalho A. B., Dobo B. A., Vibranovski M. D., Clark A. G., 2001 Identification of five new genes on the Y chromosome of *Drosophila melanogaster*. Proc. Natl. Acad. Sci. USA 98:13225–13230.

Charlesworth B., 1991 Evolution of sex chromosomes. Science 251: 1030–1033.

Charlesworth D., Charlesworth B., Marais G., 2005 Steps in the evolution of heteromorphic sex chromosomes. Heredity (Edinb). 95: 118–128.

Charlesworth D., Mank J. E., 2010 The birds and the bees and the flowers and the trees: Lessons from genetic mapping of sex determination in plants and animals. Genetics 186: 9–31.

Chen X.-G., Jiang X., Gu J., Xu M., Wu Y., et al., 2015 Genome sequence of the Asian Tiger mosquito, *Aedes albopictus*, reveals insights into its biology, genetics, and evolution. Proc. Natl. Acad. Sci.: 201516410.

Clements A. N., 1992 The Biology of Mosquitoes. Chapman & Hall, London.

Craig G. B., Hickey W. A., Vandehey R. C., 1960 An inherited male-producing factor in *Aedes aegypti*. Science 132: 1887–1889.

Fauci A. S., Morens D. M., 2016 Zika Virus in the Americas — Yet Another Arbovirus Threat. N. Engl. J. Med. 374: 601–604.

Fontaine A., Filipović I., Fansiri T., Hoffmann A. A., Rašić G., et al., 2016 Cryptic genetic differentiation of the sex-determining chromosome in the mosquito Aedes aegypti. bioRxiv.

Gilles J. R. L., Schetelig M. F., Scolari F., Marec F., Capurro M. L., et al., 2014 Towards mosquito sterile insect technique programmes: exploring genetic, molecular, mechanical and behavioural methods of sex separation in mosquitoes. Acta Trop. 132: S178–187.

Hall A. B., Timoshevskiy V. A., Sharakhova M. V., Jiang X., Basu S., et al., 2014 Insights into the preservation of the homomorphic sex-determining chromosome of *Aedes aegypti* from the discovery of a male-biased gene tightly linked to the M-locus. Genome Biol. Evol. 6: 179–191.

Hall A. B., Basu S., Jiang X., Qi Y., Timoshevskiy V. A., et al., 2015 A male-determining factor in the mosquito *Aedes aegypti*. Science 348: 1268–70.

Hall A. B., Papathanos P.-A., Sharma A., Cheng C., Akbari O. S., et al., 2016 Radical remodeling of the Y chromosome in a recent radiation of malaria mosquitoes. Proc. Natl. Acad. Sci. 113: 201525164.

Hoang K. P., Teo T. M., Ho T. X., Le V. S., 2016 Mechanisms of sex determination and transmission ratio distortion in *Aedes aegypti*. Parasit. Vectors 9: 49.

Kaiser V. B., Bachtrog D., 2010 Evolution of sex chromosomes in insects. Annu. Rev. Genet. 44: 91–112.

Koren S., Phillippy A. M., 2015 One chromosome, one contig: complete microbial genomes from long-read sequencing and assembly. Curr. Opin. Microbiol. 23: 110–120.

Laughlin C. A., Morens D. M., Cassetti M. C., Costero-Saint Denis A., San Martin J. L., et al., 2012 Dengue research opportunities in the Americas. J. Infect. Dis. 206: 1121–1127.

Matthews B. J., McBride C. S., DeGennaro M., Despo O., Vosshall L. B., 2016 The neurotranscriptome of the *Aedes aegypti* mosquito. BMC Genomics 17: 32.

Muller H. J., 1964 The relation of recombination to mutational advance. Mutat. Res. 1: 2–9.

Musso D., Cao-Lormeau V. M., Gubler D. J., 2015 Zika virus: following the path of dengue and chikungunya? Lancet 386: 243–244.

Nene V., Wortman J. R., Lawson D., Haas B., Kodira C., et al., 2007 Genome sequence of *Aedes aegypti*, a major arbovirus vector. Science 316: 1718–1723.

Newton M. E., Wood R. J., Southern D. I., 1978 Cytological mapping of the M and D loci in the mosquito, Aedes aegypti (L.). Genetica 48: 137–143.

Severson D. W., Behura S. K., 2012 Mosquito genomics: progress and challenges. Annu. Rev. Entomol. 57: 143–166.

Toups M. A., Hahn M. W., 2010 Retrogenes reveal the direction of sex-chromosome evolution in mosquitoes. Genetics 186: 763–766.

